# Multiplexed fluorescence and scatter detection with single cell resolution using on-chip fiber optics for droplet microfluidic applications

**DOI:** 10.1101/2023.05.03.539123

**Authors:** Preksha Gupta, Ambili Mohan, Apurv Mishra, Atindra Nair, Neeladri Chowdhury, Dhanush Balekai, Kavyashree Rai, Pooja Mehta, Anil Prabhakar, Taslimarif Saiyed

## Abstract

Droplet microfluidics has emerged as a critical component of several high-throughput single cell analysis techniques in biomedical research and diagnostics. However, while there has been significant progress in the individual assays being developed, multi-parametric optical sensing of droplets and their encapsulated contents remains challenging. The current common approach of microscopy based high-speed imaging of droplets is technically complex and requires expensive instrumentation limiting their large-scale adoption. Here we have adapted the principles of the widely established technique of flow cytometry to a novel optofluidic setup – the OptiDrop platform, with on-chip detection of scatter and multiple fluorescence signals from microfluidic droplets and their contents using optical fibers. The highly customizable on-chip optical fiber-based signal detection system enables simplified, miniaturized, low-cost, multi-parametric sensing of optical signals with high sensitivity and single cell resolution within each droplet. Our work on differential expression analysis of the Major Histocompatibility Complex (MHC) protein in response to IFNγ stimulation further demonstrates the OptiDrop platforms capabilities in sensitively detecting cell surface biomarkers using fluorescently labeled antibodies. The OptiDrop platform thus combines the versatility of flow cytometry with the power of droplet microfluidics to provide wide-ranging optical sensing solutions for research and diagnostics.

## INTRODUCTION

Droplet-based microfluidic technologies are providing invaluable solutions for high-throughput single cell analysis^1,2^ at the genomic^3^, epigenomic^4^, transcriptomic^5–7^, proteomic^8–11^ and metabolomic^12^ levels with increasing applications in the fields of immunotherapies^13–15^, precision medicine^16^, regenerative medicine^17^, drug development^14,18,19^ and diagnostics^12,20^. Successful translation of several such droplet-based techniques into commercial assays (e.g. ddPCR by Biorad, Chromium by 10X Genomics) has led to an unprecedented adoption of microfluidic technologies in the commercial sector. The power of droplet microfluidics stems in part from the ability to compartmentalize a larger fluid volume and its contents into much smaller nanolitre sized water-in-oil emulsion droplets. The resultant droplets act as microreactors for the rapid analysis of encapsulated single cells, and can be used for complex multistep biological, chemical assays with considerably lower reagent consumption and much faster sampling rates. Miniaturization of reaction volumes lowers the overall cost of the assays and enables rapid detection of secreted biomolecules from encapsulated cells as they quickly reach detectable levels within the smaller droplet volume^1,13^. Furthermore, droplets can be flexibly manipulated to split, merge, incorporate new reagents within preformed droplets, selectively sort out sub-populations of interest and broken down to recover viable cells^1,13,21–23^.

One key component of droplet-based assays is the detection and analysis of droplet contents. Despite the development of increasingly complex applications of droplet-based assays, there has been very limited progress in the sensing of the cells or molecules secreted by the cells trapped inside the droplets. Fluorescence readouts from droplets remains the most common approach for assessing droplet contents^24^. Traditional microfluidic setups depend on fluorescence microscope associated high speed camera imaging of droplets^13,22,23,25,26^. This off-chip optical setup often involves a complex integration and alignment of conventional free-space optics with arrays of lenses and dichroic mirrors. The entire system is rather expensive, bulky, requires skilled handling and maintenance, has limited flexibility for assay design due to fixed optical components and is restricted in terms of the number of optical parameters that can be measured simultaneously^24^. The cost and technical complexity of these optical setups have created a high-entry barrier limiting the adoption of droplet microfluidic technologies for conventional biological research and their applications in point-of-care diagnostics.

Besides fluorescence microscopy, flow cytometry has been the technology of choice for high-throughput optical analysis of single cells in conventional research and diagnostic setups^27^. In flow cytometry, light scatter and fluorescence emission signals generated by a beam of incident light shining on a stream of single cells, provide unparalleled multiparametric cellular analysis^28^. There have been several attempts to miniaturise and apply the principles of flow cytometry in lab-on-chip devices for the analysis of single cells flowing through an optofluidic channel^29–35^. However, the application of flow cytometry principles for analysis of single cells encapsulated in water-in-oil droplets remains challenging, in part due to the confounding optics of light passing through phases of different refractive indices along with the droplet size often being much larger than the cell within it. A few studies have demonstrated the detection of scatter, absorbance and fluorescence signals from dye solutions and cell populations encapsulated in droplets^13,22,25,36–39^. However, studies so far lack the sensitivity and the ability to detect and differentiate multiple optical signals with single cell resolution required for optimal high-throughput analysis. Attempts have also been made to study cells in droplets in a standard flow cytometer by making double emulsion droplets which can be suspended in the aqueous sheath fluid^40^. While being a powerful technique, double emulsion droplets are still challenging to work with and cannot be adapted for the versatile droplet-based workflows being used now^27^.

One approach to simplify and miniaturize the optical detection of droplets is to have on-chip optofluidics by integrating micro-optical components like lenses, waveguides and optical fibers within the chip design^29,35,41,42^. In particular, microfluidic chip integrated optical fibers have been demonstrated to function as effective waveguides for both the illumination of droplets by incident light and collection of optical signals from droplets^36–38^. Owing to their lower cost, ease of integration, compatibility with wide range of wavelengths, negligible transmission loss and flexibility of use, optical fibers can provide appropriate optical sensing solutions across diverse microfluidic devices.

Here, we demonstrate the integration of optical fibers on a microfluidic chip for multiparametric scatter and fluorescence analysis of droplet contents with single cell resolution. The droplet interrogation point on the microfluidic channel is flanked by a set of optical fibers placed at specific angles to the incident light fiber. The novel placement of fibers on the OptiDrop chip allows for the collection of light scattered both off the surface of the droplet as well as its contents. It also enables highly sensitive detection of multiple fluorescence signals emanating from both the solvent phase (dye/chemical solutions or secreted molecules) and from each cell or bead associated fluorophores. Thus, using a single laser as light source and multiple photo multiplier tubes (PMTs) for signal detection, the corresponding scatter and fluorescence signals from each droplet are recorded as electronic signal peaks. The recorded peak data can be used for real-time visualization and further data analysis. This simultaneous acquisition of scatter and multiplexed fluorescence signals from each droplet combines the flexibility of optofluidics with the versatility of flow cytometry and provides unparalleled optical detection capabilities for droplet-based assays. The use of on-chip optical fibers for light collection avoids the use of any free-space optical components, enabling miniaturization at much lower cost. The setup can be easily customized as per assay requirement for the combination of laser source and fluorophores by simply changing the set of optical filters attached to the PMTs for detecting the appropriate wavelengths of light.

For validation, we have characterized the OptiDrop platforms’ performance using industry standard dyes and fluorescently labelled intensity standard beads. We have further demonstrated its biological relevance using one of the most common techniques of cell surface biomarker detection using fluorescently tagged antibodies. The expression of the major histocompatibility complex (MHC) proteins on the cell surface in response to an immunogen, cytokine or external infection is often used to monitor the activity of an organism’s adaptive immune response^43^. Here we show that the OptiDrop platform is capable of simultaneously detecting differences in cell surface expression of the MHC I and MHC II molecules in response to Interferon Gamma (IFNγ) activation in mouse embryonic fibroblast (MEFs) cells encapsulated within droplets. To our knowledge, this represents the first demonstration of multiplexed fluorescence and scatter signal detection with single cell resolution using on-chip optofluidics.

## RESULTS

### Design and characterization of the OptiDrop platform

The OptiDrop platform comprises of a microfluidic chip, optical fibers for laser coupling and optical signal collection, photomultiplier tubes (PMTs) for optical signal detection, a pulse counter for signal acquisition along with algorithms for signal processing and analysis (Fig 1). The microfluidic device has a standard flow-focusing junction which allows generation of stable monodisperse nanolitre-sized water-in-oil droplets over extended periods of time^5,44^. For on-chip optical analysis, the central droplet flow channel has a set of coplanar grooves or fiber guide channels set at an angle of 45º to the incident laser guide channel at the point of interrogation (Fig 1 inset). Optical fibers placed inside the guide channels are used both for illumination with a 488 nm fiber coupled laser and collection of scattered and fluorescence light signals from the droplet passing through the point of interrogation. Optical signals collected via optical fibers are individually coupled into a photodetector, in this case a PMT with TTL (transistor-transistor logic) output. As each droplet flows through the incident beam of light, the corresponding PMT outputs are read over a period using a pulse counter. The acquired signals are captured and processed to simultaneously obtain the corresponding scatter and up to three fluorescence signals from each droplet and its encapsulated contents.

**Figure 1:**
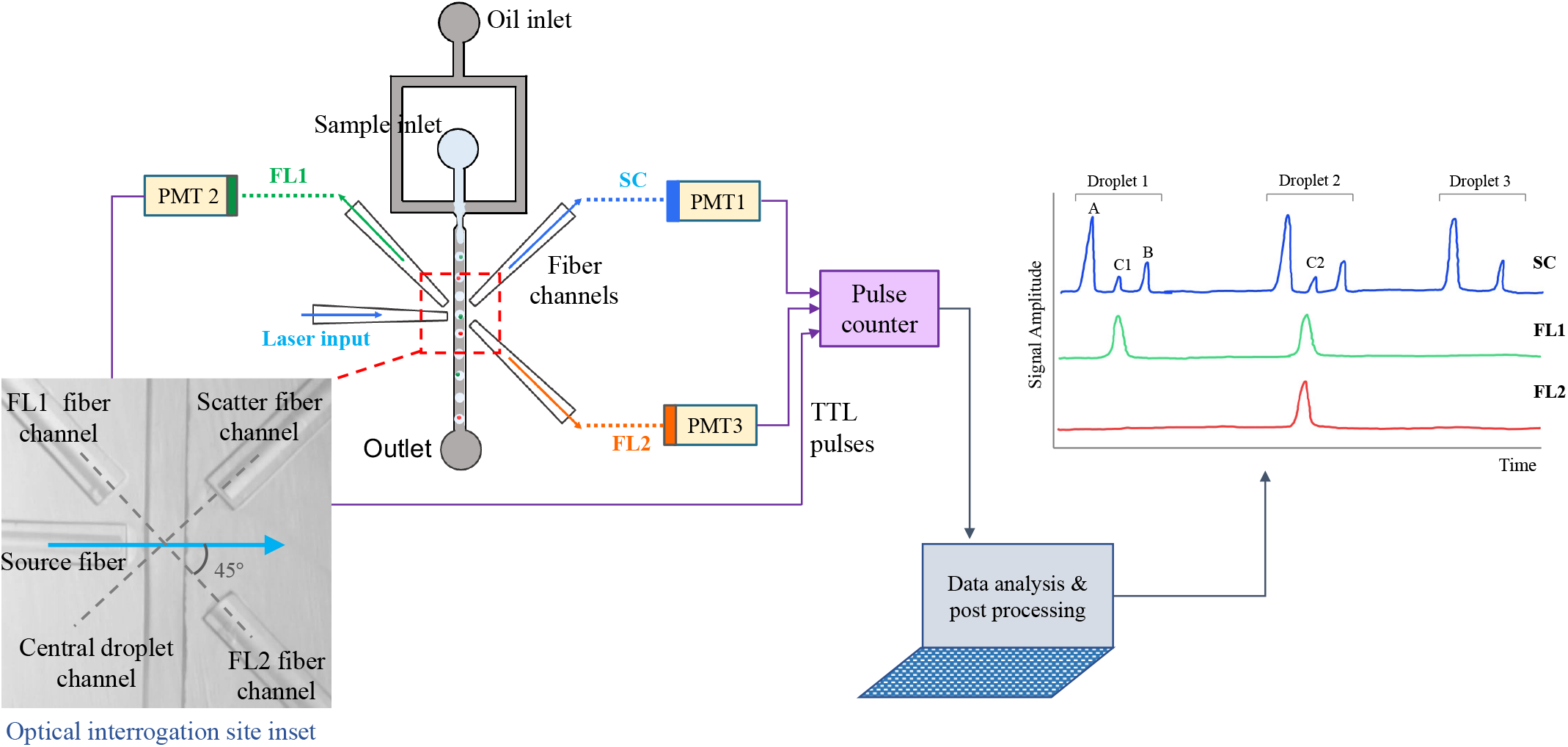
The OptiDrop platform. comprises of a microfluidic chip with an oil and water inlet for the formation of droplets at the flow-focusing junction. Contents of the aqueous phase get encapsulated within droplets and subsequently flow through the optical interrogation site (inset) flanked by a set of optical fiber grooves arranged around the central channel. The grooves on the chip are used to house the optical fibers at set angular positions allowing for effective droplet illumination with incident laser light and collection of scattered light and fluorescence signals as the droplet passes the light beam. The optical fiber output is coupled into a PMT for detection. TTL pulse counts from each PMT are integrated by an FPGA chip with a pulse counter and plotted as signal intensity peaks raw data. The raw data can be further analyzed to identify or measure fluorescence intensities from cells of interest.

#### 1. The droplet scatter signal

As each droplet travels through the incident beam, light scattered off its surface and its contents is collected via an optical fiber placed at an angle of 45º on the opposite side of the channel. The collected scatter signal is coupled into a PMT mounted with a combination of bandpass and neutral density filters. The pulse counter reads the PMT TTL pulses over a specified period and transmits the count value represented as points depicted in Figure 2. The amount of light scattered off the droplet surface is a function of the difference in refractive indices of the mediums as light passes from oil phase to aqueous phase^45^. A combination of surface reflection, refraction and total internal reflection at the droplet boundary results in the signature two peak droplet scatter signal (Fig 2B). The tall leading peak corresponds to the front of the droplet entering the incident light and the short lagging peak corresponding to the rear end of the droplet leaving the incident laser light. Thus, the leading and lagging scatter peaks effectively represent the droplet boundaries.

**Figure 2:**
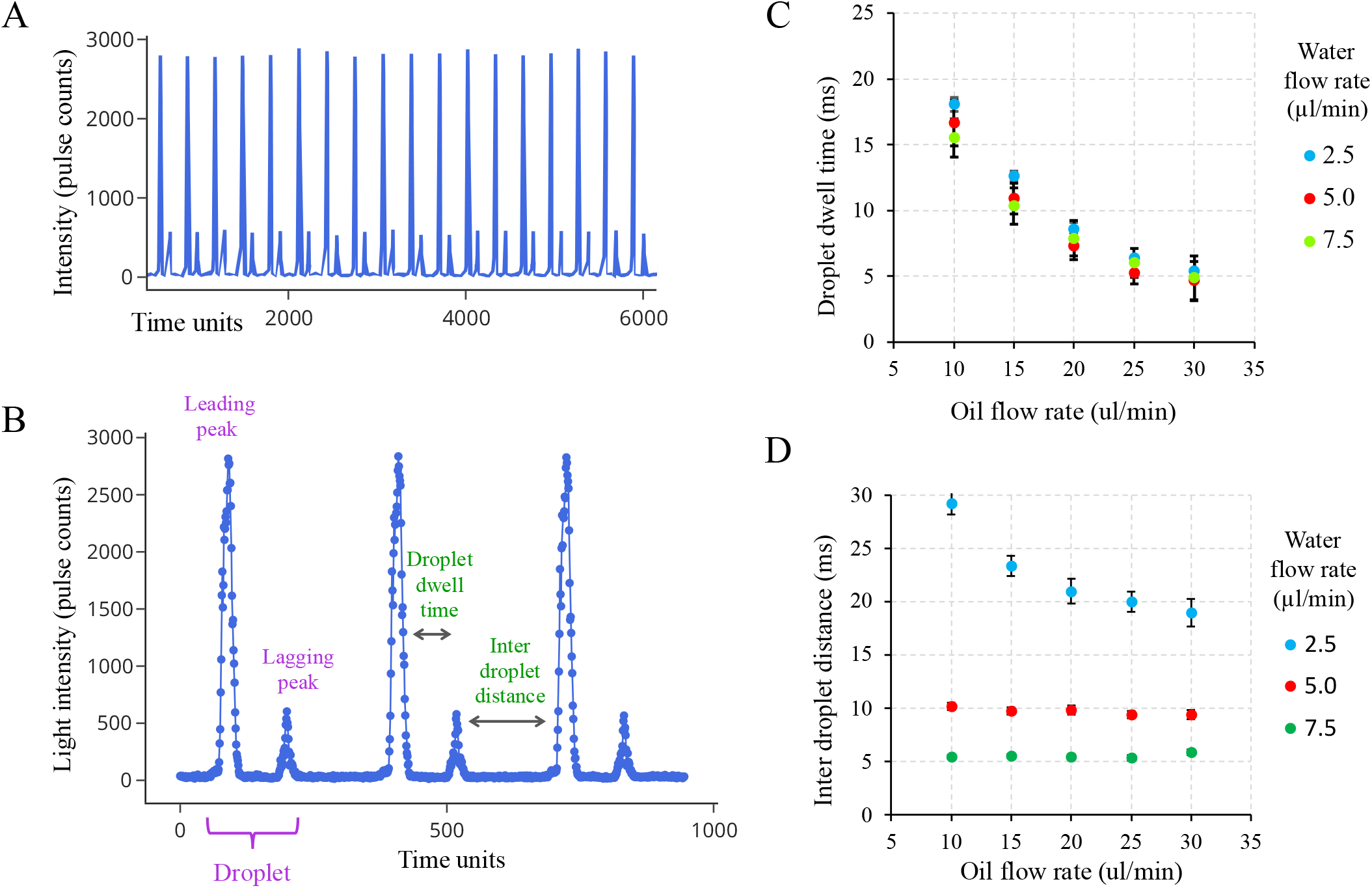
Droplet scatter signal. **A)** Raw PMT data output as count of light pulses coupled into the PMT via optical fiber over time **B)** As each droplet crosses the incident beam, it results in a characteristic two peak scatter signal of a leading and a lagging peak. Each point on the plot represents the actual pulse count from the PMT **C)** The distance between the leading and lagging peaks of the droplet or the ‘droplet dwell time’ is a function of the aqueous flow rate and independent of the oil flow rate **D)** the distance between droplets is mainly determined by the oil flow rates. Error bars indicate standard deviation.

The droplet scatter signals were further characterized by generating droplets at varying oil and water flow rates. An increase in oil flow rate while keeping the aqueous flow rate constant, results in a corresponding reduction in the intra-droplet peak distance. As oil flow rate determines the velocity of flowing droplets^46^, the faster flowing droplets spend a correspondingly lesser amount of time crossing the incident beam of light. The reduced ‘dwell time’ of the droplets, thus results in reduction in intra-droplet peak distance. Incidentally, the droplet dwell time is independent of the flow rate of the aqueous medium (Fig 2C). On the other hand, an increase in water flow rate at constant oil flow rate results in an increase in the frequency of droplet formation as evidenced by the reduction in the inter-droplet peak distance (Fig 2D).

To maximise the amount of incident light entering the droplet and the emitted light collected by optical fibers, the refractive index of oil was modified to match that of the aqueous phase^47^ by adding 3-bromobenzotrifluoride to the Novec dSURF oil. By varying the concentration of 3-bromobenzotrifluoride, we were able to carefully control the strength of droplet scatter signal as per assay requirement (Fig 3).

**Figure 3:**
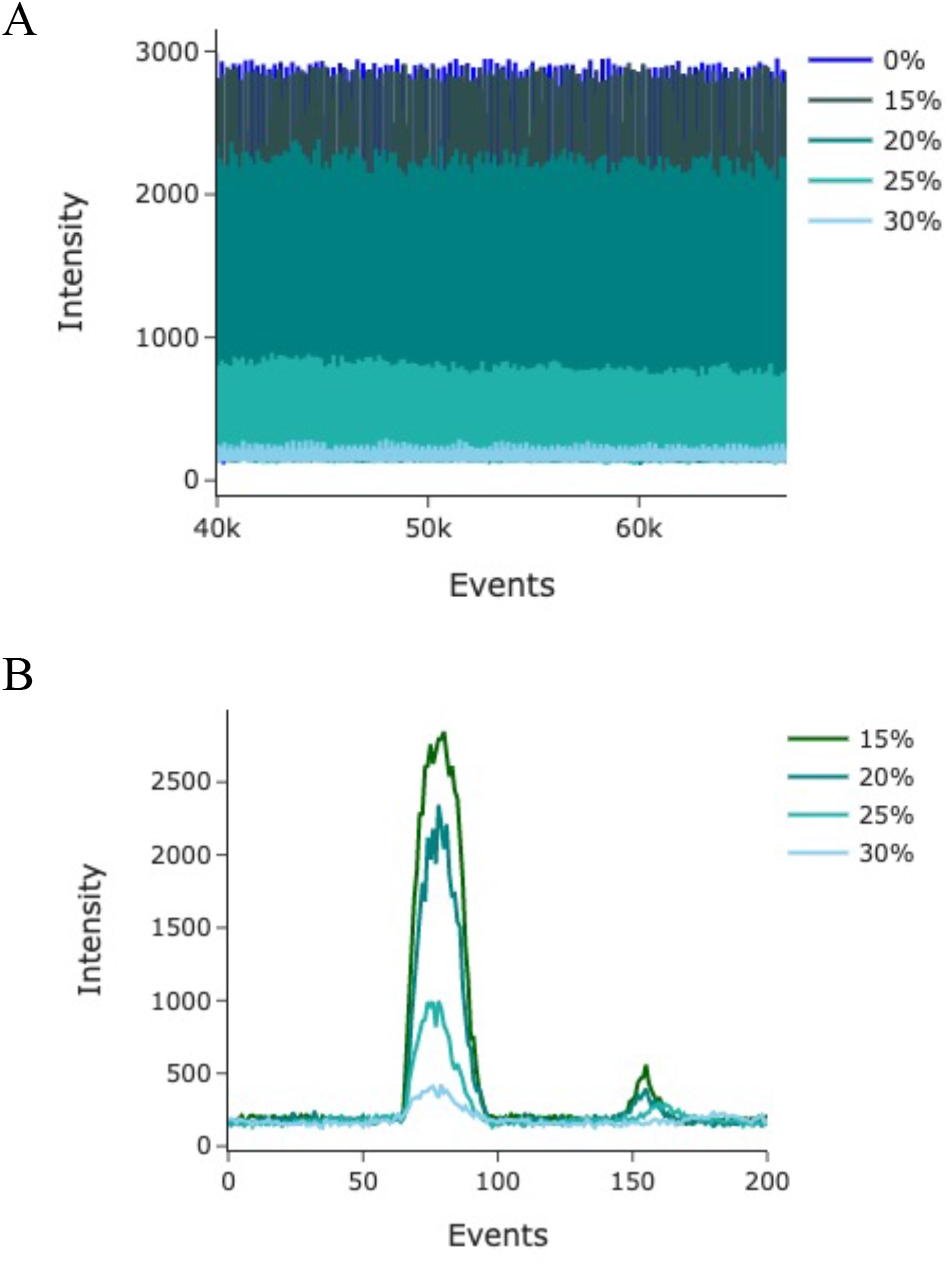
Impact of 3-bromobenzotrifluoride addition on droplet scatter signal. Increasing concentrations of 3-bromobenzotrifluoride (0-30%) when added to Novec dSURF oil, reduce the refractive index differences between the oil and aqueous phase leading to a corresponding reduction in the intensity of scatter signal as more light passes through the droplet without being refracted off its surface into the scatter collection fiber.

#### 2. The droplet fluorescence signal

Fluorescence readouts with high sensitivity are a crucial requirement of several chemical and biological assays. The OptiDrop platform can be used to collect as many as four fluorescence signals emitted from a single droplet using optical fibers placed at 45º angles to the laser guide channel (Fig 1). Light collected through each optical fiber is coupled into a separate PMT with a bandpass filter attached for the corresponding color signal being recorded. To characterize the fluorescence sensing capabilities of the OptiDrop platform, we first generated dye droplets by using increasing concentrations of the Rhodamine 123 dye as the aqueous phase and measured fluorescence intensities for each sample (Fig 4A). At a fixed oil and aqueous flow rate, uniformly formed droplets have a fluorescence signal of constant peak width and increasing peak height corresponding to the increase in dye concentration in different samples (Fig 4B). At a fixed dye concentration, the droplet fluorescence peak height remains constant while the peak width increases with increasing droplet dwell time at slower oil flow rates (Fig 4C).

**Figure 4:**
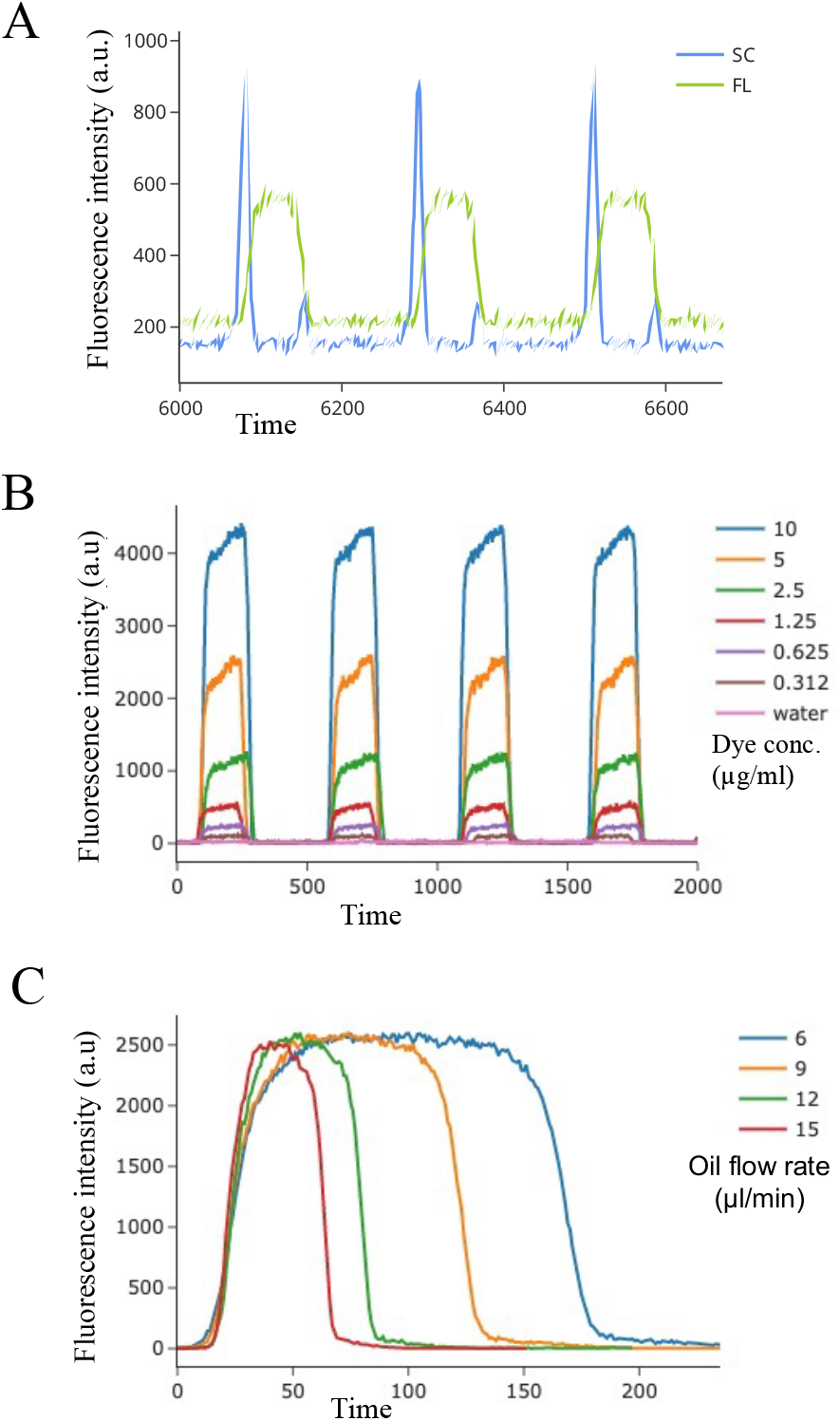
Fluorescence signal characterization of dye droplets. **A)** A typical droplet scatter (SC) and fluorescence (FL) signal acquisition from droplets encapsulating fluorescent dye solutions. **B)** Overlap plot of droplet fluorescence signals from droplets encapsulating increasing concentrations of Rhodamine 123 dye (0-10 μg/ml) solutions recorded at the same flow rates. **C)** Overlap plot of droplet fluorescent signals acquired from droplets encapsulating the same Rhodamine 123 dye solution at varying oil flow rates.

We then examined the linearity and dynamic range of detection of the OptiDrop platform. Increasing dye concentrations of fluorescein resulted in a linear increase (R^2^ value of 0.99) in the droplet fluorescence intensities with a high dynamic range of detection spanning from 1 nM to 1 μM for 4 nL droplets (Fig 5). To determine the detection efficiency of the OptiDrop platform, we tested the limit of detection (LOD) for fluorescein (an industry reference standard) and phycoerythrin dye droplets. The LOD, defined as the concentration which gives a signal equal to three times the standard deviation of the blank measurements, was determined to be 1 nM of fluorescein (Fig 5). Similarly, LOD for phycoerythrin was found to be as low as 0.5 pM (Fig S1). Fluorescence signals were also measured on a standard benchtop fluorimeter (Varioscan Lux, Thermo Fisher Scientific) by loading 20 μl of the dye solutions in a 384 well plate for comparison (Fig S2). These results demonstrate the capabilities of the OptiDrop platform in sensitively detecting multiple fluorescence signals from within droplets with LODs comparable to or better than those reported previously in similar optofluidic devices ^36,38^. Here we have shown representative data for droplets encapsulating single dye solutions, but the setup can further easily be used to detect fluorescence signals from a mixture of dyes encapsulated within droplets by collecting optical signals simultaneously at different angles.

**Figure 5:**
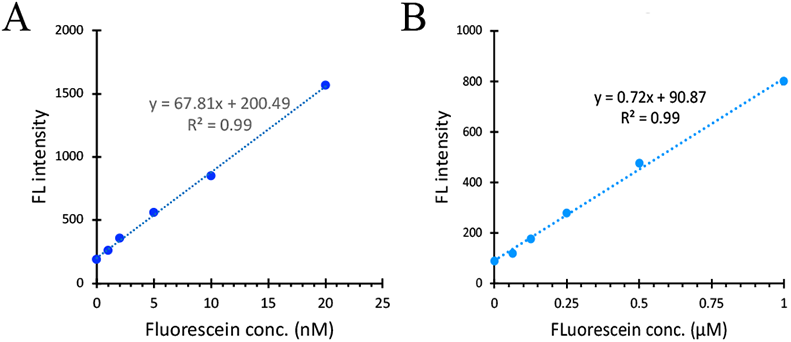
Detection efficiency for Fluorescein dye. Fluorescence signal intensity shows a linear increase with increasing fluorescein dye concentrations and has a high dynamic range A) 0-20 nM B) 0-1 μM concentrations. The LOD for fluorescein is determined ot be 1 nM.

#### 3. Encapsulated single cell/bead scatter and fluorescence signals

The OptiDrop platform provides not only whole droplet scatter and fluorescence signals, but also multiparametric optical information from each droplet constituent cell/bead with single-cell resolution. During a typical run, the scatter and corresponding fluorescence signals from each cell/bead are captured as individual peaks within the droplet boundary scatter peaks (Fig 6A). For instance, in a sample of fluorescent microbeads randomly distributed in droplets following Poisson distribution, the OptiDrop platform can effectively distinguish between droplets containing zero, one or two beads. A double bead containing droplet has 2 peaks each corresponding to the bead scatter and fluorescence signals present within the leading and lagging droplet scatter peaks (Fig 6A).

**Figure 6:**
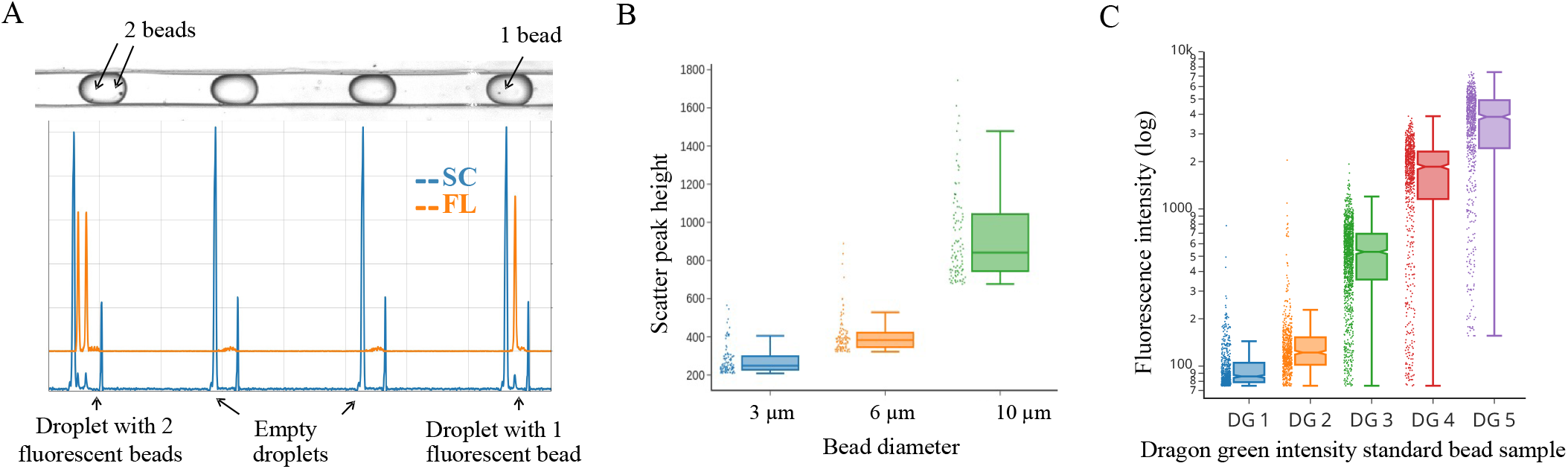
Scatter and fluorescence signals from beads encapsulated within droplets. **A)** Snapshot of the droplets flowing through the microfluidic channel (top image) and its corresponding data recorded on the OptiDrop platform (below plot). 4 droplet boundaries are demarcated by the leading and lagging scatter peaks. Within the boundary peaks lie the bead scatter and fluorescence peak. Scatter (SC) signal in blue and fluorescence (FL) signal in orange. **B)** Box plot indicating the size distribution of scatter signal from 3, 6 and 10 μm sized beads. **C)** Box plot demonstrating the increasing fluorescence intensity profiles of the five populations of the Dragon Green intensity standard kit.

The height of the scatter peaks roughly corresponds to the bead size. An increase in the size of the 3, 6 and 10 μm sized polystyrene particles encapsulated in droplets, results in a corresponding increase in their scatter signal intensity (Fig 6B). Similarly, the height of the fluorescence peaks corresponds to the signal intensity associated with each cell/bead. To characterize the platform’s ability in measuring relative fluorescence intensities, we used the Dragon Green intensity standard kit (Bangs Laboratories, Inc). The OptiDrop platform can sensitively distinguish between the five populations of polystyrene microbeads dyed with increasing amounts of Dragon Green fluorophore (Fig 6C).

In order to understand the limit of detection for bead or cell associated fluorescence, we used Rainbow flow cytometry calibration particles (Spherotech) as recommended standards for biological applications. Peak 2 bead population corresponds to the dim beads often used in standard flow cytometers to set the instrument for detecting the lowest fluorescence signal measured above the blank signal while the Peak 5 population corresponds to the brighter end of the spectrum of bead fluorescence. Fluorescent signals from both the dim peak 2 and the bright peak 5 population of beads were effectively captured (Fig S3).

To validate the robustness of the OptiDrop platform, 80% FITC stained CaliBRITE beads were mixed with 20% PE stained CaliBRITE beads. FITC and PE Fluorescent signals were acquired from the mixed bead sample encapsulated within droplets and also on a standard flow cytometer for comparison. The distribution of FITC and PE beads on the scatter plot recorded on the OptiDrop platform in Fig 7 shows the expected 8:2 signal ratio. Similar results are observed on a standard flow cytometer (Fig S4). This experiment demonstrates the platforms capability in effectively capturing fluorescence read outs from each bead in the population.

**Figure 7:**
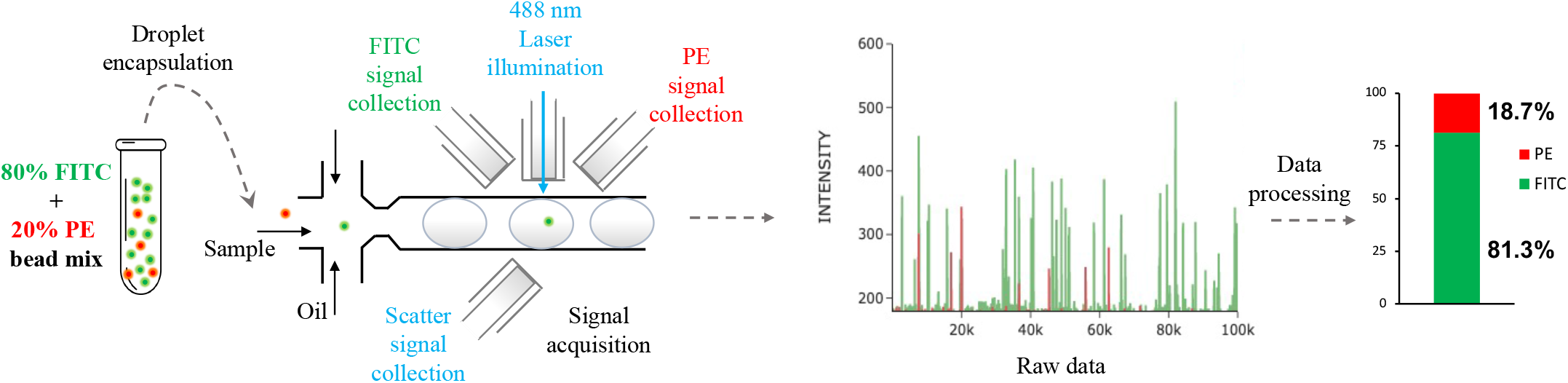
Effective signal acquisition and differentiation in a mixed bead population. A mixture of FITC (80%) and PE (20%) stained CaliBRITE bead sample data is anlyzed on the OptiDrop platform. The acquired raw fluorescence peak signals are analyzed and subsequent the proportion of FITC and PE beads is calculated as shown in the bar plot.

#### 4. Biological application: MHC cell surface expression profiling in MEFs

Antibody-based protein biomarker detection is the most common method for assessing the expression of cell surface proteins using flow cytometry with versatile applications in both biomedical research and diagnostics. One such application is the analysis of MHC immunomodulation in cells. MHC proteins play a critical role in the immune system by presenting antigens to T cells. MHC molecules are divided into two main classes: MHC class I and MHC class II molecules. MHC class I molecules are expressed on the surface of most nucleated cells and present antigenic peptides derived from intracellular pathogens, such as viruses or intracellular bacteria, to CD8+ cytotoxic T cells. MHC class II molecules, on the other hand, are primarily expressed on the surface of antigen-presenting cells, such as dendritic cells, macrophages, and B cells. These molecules present antigenic peptides derived from extracellular pathogens, such as bacteria or parasites, to CD4+ helper T cells. Together the MHC class I and II molecules enable the detection and elimination of infected cells by activating the appropriate immune response^48^. Expression levels of MHC molecules on the surface of cells, can thus serve as an important indicator of an organism’s ability to fend off pathogens. Recent studies have suggested that specific MHC alleles and their expression may be associated with increased susceptibility to COVID-19 infection or more severe disease outcome^49,50^. Additionally, abnormalities in MHC expression can be used as a diagnostic tool for various diseases, including autoimmune disorders and cancer^51,52^.

Given the importance of MHC proteins in understanding immune system function and developing therapeutic strategies for immune-related diseases, we attempted to analyze their expression in mouse embryonic fibroblasts (MEFs) in response to IFNγ activation. Utilizing the OptiDrop platform’s ability to simultaneously detect scatter and two fluorescence signals from each encapsulated cell, we used PE and FITC conjugated antibodies to stain cell surface MHC class I and class II proteins respectively. Stained cells were encapsulated in droplets and multiparametric analysis was performed on the OptiDrop platform. Unstimulated MEFs have a low basal expression of MHC class I proteins and do not express any MHC Class II molecules. The OptiDrop platform was effectively able to sense the increase in MHC class I proteins and the activation of MHC class II expression in response to IFNγ stimulation (Fig 8). The results obtained on the OptiDrop platform were validated to be comparable to that obtained on a standard flow cytometer (Fig S5).

**Figure 8:**
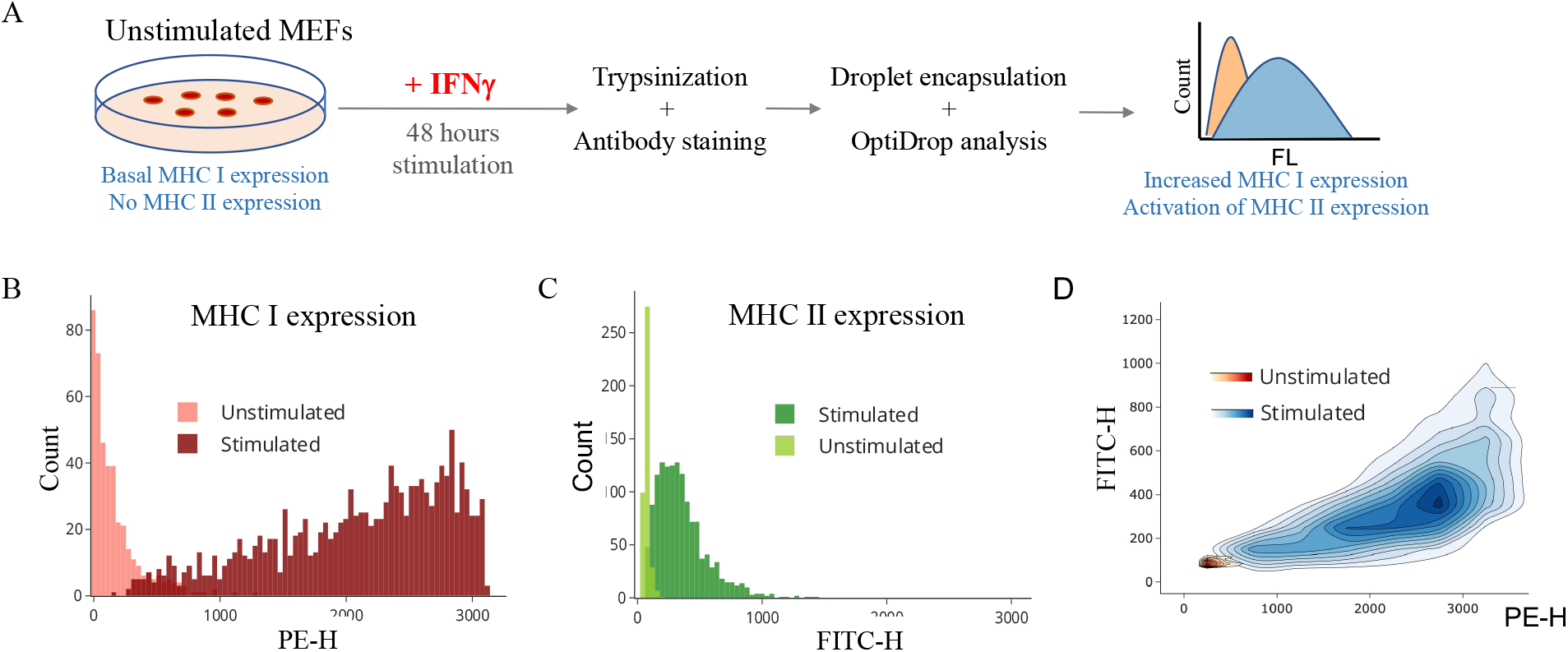
MHC protein expression profiling in response to IFNγ stimulation. **A)** Schematic of the biological assay depicting the stimulation of MEFs with IFNγ for 48 hours, subsequent removal of cells from culture plate using Trypsin, staining with PE-tagged MHC I and FITC-tagged MHC II antibodies. Stained cells were encapsulated within droplets on the OptiDrop platform and the fluorescence signal was acquired. **B), C)** The fluorescence intensities from each cell of stimulated and unstimulated MEF samples is plotted as histograms depicting MHC I and MHC II expression. D) Two color fluorescence density contour plot depicting the relative increase in both MHC I and MHC II expression upon stimulation with IFNγ.

## MATERIALS AND METHODS

### A. Device fabrication protocol

The microfluidic chips were fabricated using polydimethylsiloxane (PDMS) by standard soft-lithography techniques as previously described^29^. Briefly, the design mold was fabricated on a silicon wafer by UV exposure through a photo-lithography mask and subsequent development. A curing agent was added to the PDMS base (Sylgard 184 silicone elastomer kit; Dow Corning Corporation) to a final concentration of 10 % by weight, mixed and poured over the mold to a depth of 7 mm. Following degassing for several minutes and cross-linking at 70ºC for 2.5 hours, the polymerized PDMS was peeled off the mold and cut into individual chips. The inlet and outlet ports were punched in the PDMS with a 0.9 mm diameter biopsy punch for fluidic connections.

PDMS layer was subjected to the oxygen plasma treatment (Harrick plasma cleaner) with patterned side facing up and the substrate layer (cleaned glass slides) adjacent to it. Vacuum was created in the plasma chamber for 45 seconds. Oxygen was slowly purged into the plasma chamber until the chamber color turned dark with a pink periphery. The oxygen purge was stopped upon the color change and the chamber left undisturbed for a minute or two before the vacuum was released to open the chamber. After taking them out from the chamber, the PDMS device was permanently bonded onto the glass slide by pressing it against the glass slide without excessive pressure. The bonded device was further placed in a vacuum oven at 70ºC for 25 to 30 minutes. The central fluidic channels were treated with Aquapel^5^ and kept in hot air oven at 70ºC for 45 minutes before using.

### B. Device setup

The sample containing any cells, beads or dye solutions comprised the aqueous phase. Fluigent dSURF oil (2% surfactant in 3M™ Novec™ 7500 fluorinated oil) was used as the continuous phase. The sample and oil inlets were connected to the syringe pumps (NE 300, New era pump systems Inc.) via polyethylene tubing (BTPE-50, Instech laboratories Inc.) for fluid injection. The flow-focusing junction on the microfluidic chip breaks the flow of aqueous sample to form water-in-oil droplets with the contents of the aqueous sample getting encapsulated within droplets. The frequency, speed and size of droplets can be controlled by modulating the oil and aqueous sample flow rates in the syringe pump.

Downstream of the flow-focusing junction, the microfluidic chip has grooves on both sides of the central flow channel for housing optical fibers (Fig.1). Lensed optical fibers (multimode 105 μm core/125 μm cladding with a working distance of 150 μm from Lase Optics Corporation) or non-lensed fibers (standard multimode OM1 fibers with 62.5 μm core/125 μm cladding diameter) are used as excitation fibers for laser light coupling. Non-lensed OM1 fibers are used for collecting scattered light and fluorescent signals from the droplets. The tips of non-lensed optical fibers were stripped off their outer protective jacket layer with only the core and cladding layer remaining. Stripped fibers were cleaved with a fiber cleaver to obtain a flat surface at the fiber tip end.

The fiber guide channels/grooves were filled with index matching fluid (Eagle Photonics). Optical fibers were then manually inserted into the guide channels with careful observation under a stereo microscope ensuring the absence of any air bubbles between the fiber tip and the end of fiber guide channel. The chip was placed on a holder and the screws tightened to secure the fibers in place. The collected light signals from optical fibers are coupled into PMT detectors (Hamamatsu H10682, Sens-Tech P25PC). Neutral density and bandpass filters are placed between the fiber and PMT interface as per the wavelengths of interest and laser intensity.

### C. Data collection and signal processing

TTL-level voltage pulses from the PMTs were processed by implementing a pulse counter for each of them using the FPGA (Field Programmable Gate Array) chip on the Digilent Zedboard (Xilinx Zynq 7000 SoC). Verilog and the Xilinx Vivado suite were used to program the hardware on the FPGA to count the pulses and store them in memory (BRAM). The integration time for pulse counting was set to 100μs within the hardware. The counts from each PMT were read simultaneously from the BRAM using interrupts, then combined into a single ethernet frame and sent out in a continuous data stream, using C and XIlinx SDK libraries on the ARM chip.

A Python and Qt based GUI was created to stream the live data output and to store the data. PMT outputs are overlaid on the GUI to obtain the corresponding scatter and fluorescence values at each timestamp.

PMT pulse count data is processed to identify peaks using our custom Python based signal extraction algorithm. Leading and lagging peaks in the scatter signals are used to mark the droplet boundaries. Within each droplet boundary, individual peaks for cell/bead scatter and fluorescence are identified and assigned to the respective droplet. Each peak data is analyzed to obtain the corresponding peak height (signal intensity) values.

### D. Chemicals and reagents

Fluorescein (Sodium salt SRL 55091) and R-Phycoerythrin (52412 Sigma Aldrich) was used for making dilutions for determining the lower limit of detection for the setup. 3-bromobenzotrifluoride (Sigma B59004) was mixed with dSURF oil for refractive index modifications as per requirement. Dragon Green intensity standard kit (DG06M) was purchased from Bangs Laboratories, Inc. Rainbow Calibration particles (RCP-60-5-2, RCP-60-5-5) were obtained from Spherotech. FITC and PE stained beads were part of the three color BD CaliBRITE kit (BD Biosciences 340486). Beckman Coulter CytoFLEX Daily QC Fluorospheres (B53230), Spherotech SPHERO Rainbow Calibration Particles, Peak 2 (RCP-60-50-2) and Polystyrene DVB-COOH microspheres from Bangs Laboratories, Inc (PC07001) were used as reference standards for 3, 6, 10 μm sized beads respectively.

### E. Cells and culture conditions

C57Bl/6 embryos were eviscerated at day 12.5-13.5 and embryonic fibroblasts were isolated at the Stem Cell Core (InStem, Bangalore). Cells were cultured in DMEM (Gibco) supplemented with 10% FBS at 37°C with 5% carbon dioxide. Cells were grown till 70-80% confluency and then treated for 48 hours with 1 μM recombinant mouse interferon gamma (Sino Biological, China). Treated cells were detached by trypsinisation and stained with antibodies against the MHC cell surface proteins.

Briefly, 1×10^6^ cells were transferred into 1.5 mL microcentrifuge tubes (Eppendorf, Germany) and washed twice with ice cold FACS buffer containing 5% FBS in PBS. Cells were centrifuged at 1000 rpm for 5 minutes in a MiniSpin Plus microcentrifuge (Eppendorf, Germany). Washed cells were resuspended in 150 μL of FACS buffer and treated with anti-mouse MHC class I antibody conjugated to PE (Clone:28-14-8) (Invitrogen, USA) at a concentration of 2μg per sample and anti-mouse MHC class II antibody conjugated to FITC (Clone: M5/114.15.2) (Invitrogen, USA) at a concentration of 5μg per sample. Cells were stained for 1 hour at a temperature of 4°C with shaking at 600 rpm in a Thermomixer comfort (Eppendorf, Germany). Stained cells were washed with ice cold FACS buffer twice before resuspension in FACS buffer for analysis. 15% OptiPrep (D1556 Sigma Aldrich) was added to the aqueous sample to prevent settling of cells within droplets.

## CONCLUSIONS & DISCUSSION

We have developed a novel platform for simultaneous scatter and multiplexed fluorescence signal detection from solutions and cells/beads encapsulated within microfluidic droplets with single cell resolution. The OptiDrop platform is sensitive (1nM fluorescence detection limit), has a high dynamic range of detection for both solution and cell/bead-based fluorescence, and is capable of differentiating cells/beads of different sizes based on their scatter signal. We believe it to be the first non-imaging platform capable of integrating the principles of flow cytometry for high throughput multiparametric single cell analysis in microfluidic droplets.

The OptiDrop platform has several advantages over the existing droplet optical analysis platforms. First, the simplified on-chip fiber optics overcomes the limitations of cost, complexity, bulkiness, and lack of flexibility associated with traditional methods of fluorescence microscopy based high-speed video imaging. The replacement of expensive, bulkier free-space optical components with multimode optical fibers enables miniaturization, significantly reduces cost, circumvents the need for complex alignments and maintenance while making it easily customizable based on assay requirements. Moreover, imaging of high-throughput droplet-based assays results in large datasets requiring expensive storage and processing solutions, further limiting their scalability. The live data visualization, small data storage footprint and simpler data analysis algorithm incorporated within the OptiDrop platform further enhances its applicability as a benchtop clinical technology.

Similarly, while the current commercial flow cytometers are highly proficient and versatile in their biomedical applications, their applicability is limited when it comes to analysing cells encapsulated within water-in-oil droplets. Also, flow cytometers are often only used for end-point analysis precluding any dynamic monitoring of single cell responses over time. The most important distinction of flow cytometers from the OptiDrop platform, in fact stems from the ability of microfluidic droplets to act as microchambers for single cells or organoids and their extracellular secretions. Droplet-based assays, and thus in essence, the OptiDrop platform, can not only provide information from single cells but can also detect secreted molecules encapsulated within each respective droplet. Another critical feature of the OptiDrop platform is its ‘closed’ nature – the biological sample remains within easily replaceable tubings, microfluidic cartridges and never goes through a system open to external contamination. Such a closed optical flow detection system, as opposed to the open system of standard and imaging flow cytometers like ImageStream, is particularly suited for clinical applications.

Although here we have validated the effectiveness of the OptiDrop platform for profiling cell surface protein expression using antibodies, its simplicity allows for seamless integration with the multiverse of existing droplet microfluidic workflows for optical sensing. Current single-cell genomic technologies provide a high-resolution view of gene expression and other genomic features, but fail to capture other aspects of cellular behaviour, such as protein expression or cell morphology. Integrating single-cell genomic data with cellular phenotype information obtained by concurrent multiparametric on-chip optical analysis using the OptiDrop platform can provide a more comprehensive understanding of cellular behaviour. The reduced cost and complexity of optical analysis can further boost the adoption of droplet microfluidics for point-of-care diagnostics.

The current work lays the foundation for further improvements in speed, sensitivity, accuracy, and cost of optical sensing of single cells encapsulated in droplets. We hope the range of applications of the OptiDrop platform technology will be extended further and that it enables the development of affordable, scalable benchtop droplet microfluidic solutions in biomedical research and diagnostics.

