## Supplementary figures for "Multiplexed fluorescence and scatter detection with single cell resolution using on-chip fiber optics for droplet microfluidic applications"

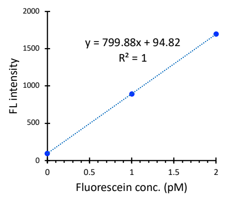


Figure S1: Standard curve for increasing concentrations of Phycoerythrin dye in droplets results in linear increase in fluorescence intensity with an LOD of 1 pM

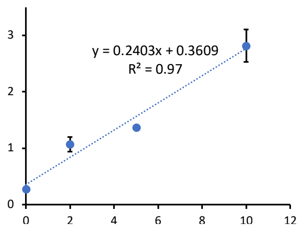


Figure S2: Fluorescence intensity measurement of 20ul Fluorescein dye solutions of increasing concentrations on a standard fluorimeter using a 384 well microplate – Varioscan LUX from Thermo Fisher. LOD was observed to be 2nM. Larger margin of eror as indicated by error bars depicting standard deviation on the linear equation with a $R^2$ value of 0.97.

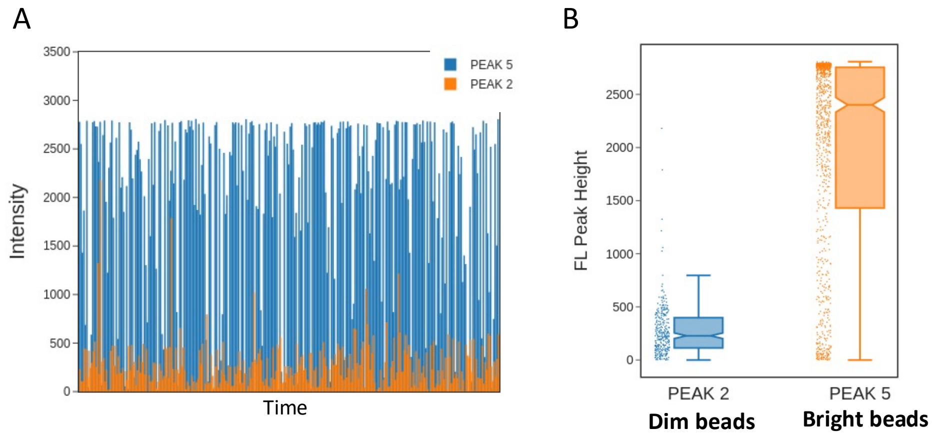


Figure S3: Fluorescence intensity measurements from the Peak 2 (dim) and Peak 5 (bright) populations of the Rainbow flow cytometry calibration particles (Spherotech) A) Raw data B) Box plot of fluorescence peak heights

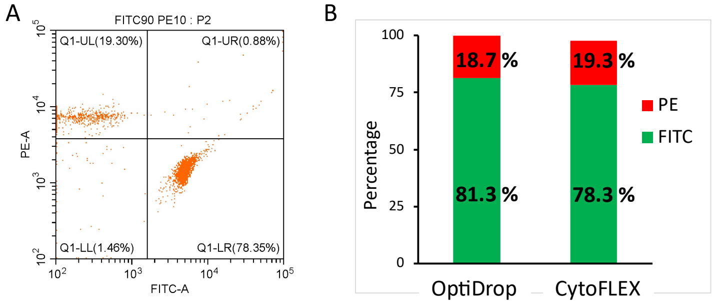


Figure S4: Comparison of OptiDrop data on mixed beads with that acquired on a standard flow cytometer – CytoFLEX.

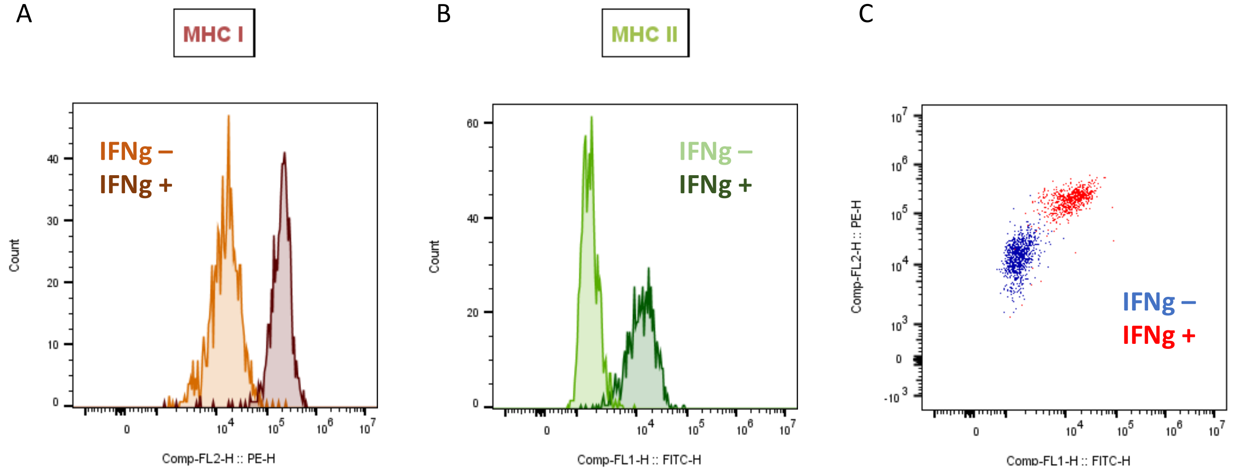


Figure S5: MEF MHC staining data obtained on standard flow cytometer (Cytoflex) is comparable to that obtained on OptiDrop platform.
